# Microglia detection and phagocytosis of dying neurons is regulated by CX3CR1

**DOI:** 10.64898/2026.04.02.716180

**Authors:** Maryanne N. Barasa, Alicia N. Pietramale, Robert A. Hill

## Abstract

Neuronal cell death is a hallmark of many neurodegenerative diseases. Effective detection and clearance of cell debris generated during cell death events is essential to prevent a degenerative cascade. Brain resident microglia are responsible for performing these functions through complex cell-cell signaling involving both “find-me” and “eat-me” cues. To examine microglial responses to neuronal cell death in vivo, we investigated neuron/microglia CX3CL1/CX3CR1 signaling using intravital optical imaging in mouse cortex and a single-cell ablation technique called 2Phatal. We find that CX3CL1 aggregates as puncta on microglia and that this pattern is maintained when microglia engulf dying neurons. Additionally, disruption of this signaling via *Cx3cr1* deletion when both few and many neurons are dying leads to delayed cell corpse clearance, partly due to a delay in microglial engagement with the dying cells. Overall, our work uncovers a precise role for CX3CL1/CX3CR1 signaling in regulating the microglial response to dying neocortical neurons.

## INTRODUCTION

Progressive neuronal loss is the fundamental pathological feature underlying various neurodegenerative diseases such as Alzheimer’s Disease (AD) and Parkinson’s Disease (PD)^1–3^. Concomitant with widespread neuronal cell death, it is now well established that microglia, the resident immune cells of the central nervous system (CNS), play an active role in the pathogenesis of these disorders^4,5^. In their canonical role as macrophages, microglia support the CNS environment during pathology by facilitating the phagocytosis of damaged and dying neurons, however, this important function is often disrupted in these disease contexts contributing to neuroinflammation and further neuronal loss^6^. More specifically, single nucleotide mutations in key microglial receptors that mediate the detection and clearance of dying neurons have been associated with increased risk of developing both AD and PD^7,8^. Among the microglial-expressed receptors implicated in mediating neuronal cell clearance, CX3CR1 remains a focal point of research given its high expression levels on microglia and its unique signaling axis with its ligand, CX3CL1 (also called fractalkine), which is predominantly expressed by specific subsets of neurons in the CNS^9–13^.

In the healthy brain, CX3CL1 is located on the outer membrane of neurons and is constitutively cleaved, primarily by ADAM10, to facilitate homeostatic communication between neurons and microglia^14^. During pathological or other inflammatory conditions, CX3CL1 can be cleaved in stressed or dying neurons by additional proteases such as ADAM17 and released as a soluble “find-me” signal to attract microglia toward the site of injury^15,16^. However, in which specific cellular contexts CX3CL1 functions as the predominant “find-me” signal in the CNS remains unclear.

Beyond chemotaxis, the CX3CL1/CX3CR1 signaling axis also acts as a critical mediator in the phagocytic action of microglia although its exact role in this regard is debated. Physiologically, CX3CL1 signaling with CX3CR1 maintains microglia in a quiescent surveillant state while inhibiting the microglial transition to a phagocytic phenotype^17–21^. In line with this, studies using multiple AD mouse models (5xTg, R1.40, and CRND8) show that *Cx3cr1* knockout (*Cx3cr1*^-/-^) microglia have an increased phagocytic capacity where the deletion of *Cx3cr1* consistently led to a gene-dose dependent increase in the clearance of protofibrillar amyloid-beta plaques^22,23^. Conversely, studies using the 5xFAD mouse model of AD^24^ as well as multiple tau pathology models^11,25–27^ show that *Cx3cr1* deletion reduced microglial phagocytosis of amyloid-beta and pathological tau respectively, both in vitro and in vivo, all while promoting both tau hyperphosphorylation and cognitive deficits. In line with the latter studies, previous research studying microglial clearance of dying cells show that the deletion of *Cx3cr1* or its respective ligand, *Cx3cl1*, disrupted microglial phagocytic capacity leading to a delay in clearance and an accumulation of cell corpses^28–30^. These seemingly conflicting observations highlight the context dependent nature of the CX3CL1/CX3CR1 signaling axis in mediating microglial phagocytosis and clearance, suggesting differential roles in distinct pathological scenarios. This context dependence therefore presents a major bottle neck in the study of CX3CR1 signaling in microglia. Experimental approaches where we can isolate specific pathological events for instance, the programed cell death of neurons in vivo, would minimize confounding systemic and inflammatory variables and isolate the role of this signaling axis in microglial responses to neuronal cell death.

To do this, we used a non-inflammatory photochemical ablation method called 2Phatal^31^ to induce apoptosis in individual neurons in conjunction with longitudinal intravital 2-photon imaging to determine if CX3CR1 signaling is important for the clearance of dying neurons. Our combined strategy not only enabled us to track precise microglial responses to dying neurons with high spatiotemporal resolution in the live brain but also allowed us to control the number of neurons undergoing cell death. Overall, this work uncovered a precise role for CX3CL1/CX3CR1 signaling in the microglial clearance of single-dying neurons in vivo and highlights this axis as an important pathway involved in brain homeostasis and neurodegenerative conditions.

## RESULTS

### CX3CL1 localizes to neurons and as puncta on microglia

Previous studies have demonstrated that in the mouse brain CX3CL1 expression is restricted to neurons^10^. To characterize the localization of CX3CL1 in the healthy mouse brain, we immunostained for CX3CL1 and the neuronal marker, NeuN (Figure 1a). As expected, CX3CL1 labeled neuronal somas (Figure 1a). CX3CL1 immunoreactivity was highest in the regions with the greatest neuronal density namely, layers II-VI, whereas layer I showed minimal CX3CL1 immunoreactivity and contained few neuronal somas. Similarly, the corpus callosum white matter tract exhibited few to no neuronal somas with little to no CX3CL1 immunoreactivity^32^ (Supplementary Figure 1). Interestingly, CX3CL1 labeling across neurons was heterogenous with some neurons exhibiting strong perineuronal/membrane associated CX3CL1 signals, whereas others showed minimal to undetectable signals (Figure 1a).

**Figure 1:**
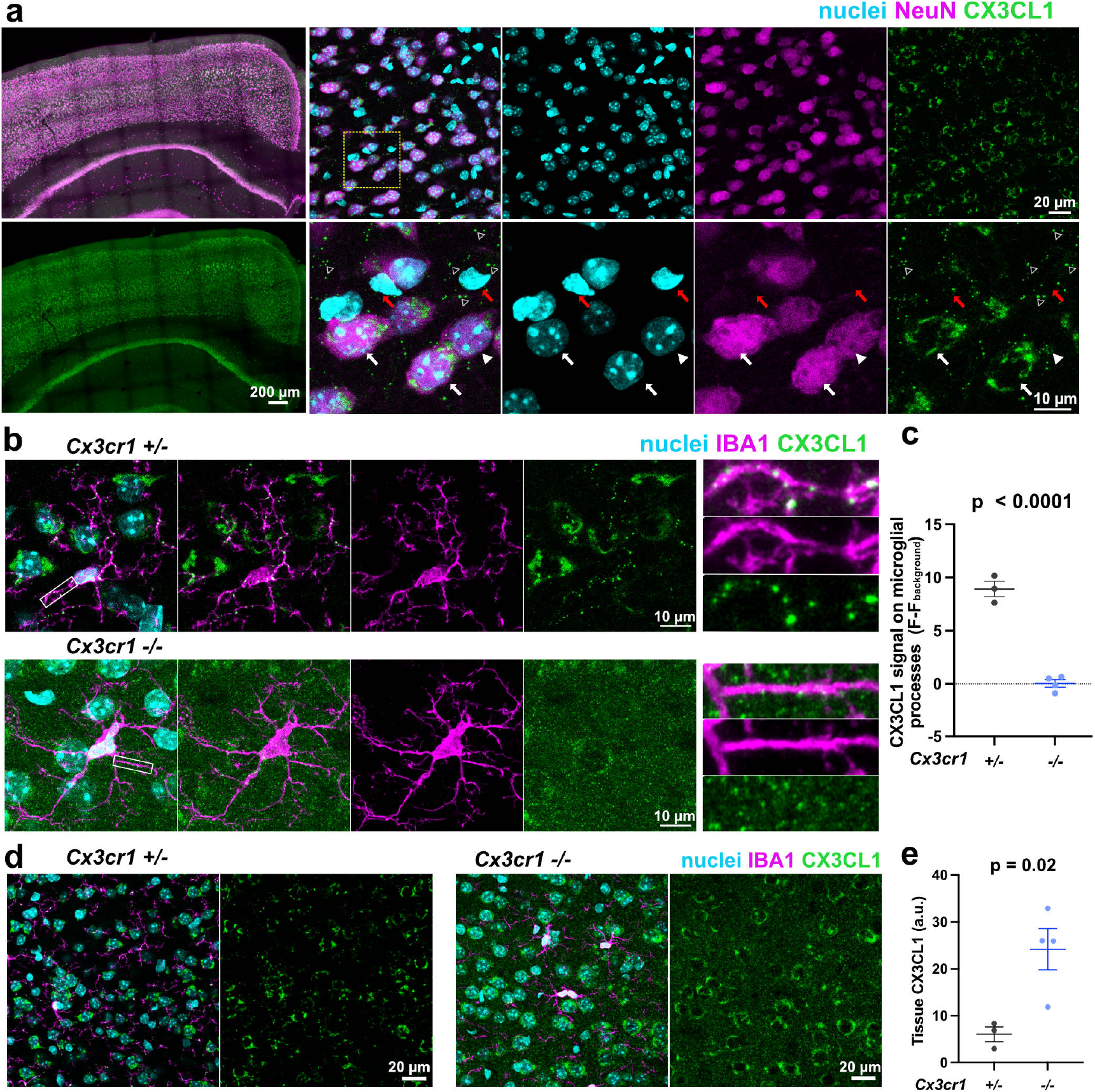
Neurons release CX3CL1 that binds CX3CR1 on microglia. **a**. Representative images of mouse coronal brain sections co-immunolabeled with antibodies against NeuN and CX3CL1. Cell nuclei are labeled with Hoechst dye. Left panels show images of fractalkine overlapping with NeuN specifically on neuronal somas. Red arrows show NeuN^-^ CX3CL1^-^ glial nuclei. White arrows show NeuN^+^ neuronal nuclei with high CX3CL1 immunoreactivity. White arrowheads show a NeuN^+^ neuronal nucleus with low CX3CL1 immunoreactivity. Gray arrowheads show fractalkine puncta not associated with neuronal nuclei. **b**. Outside of neuronal labeling, CX3CL1 puncta are present on the cell membrane of IBA1^+^ *Cx3cr1*^+/-^ microglia but are absent on IBA1^+^ *Cx3cr1*^-/-^ microglia. **c**. The CX3CL1 signal on microglial processes in *Cx3cr1*^+/-^ and *Cx3cr1*^-/-^ mice (n = 3 *Cx3cr1*^+/-^ mice and 4 *Cx3cr1*^-/-^ mice; Forty-eight processes were analyzed per genotype; Unpaired two-tailed t-test; Error bars represent SEM). **d**. Representative images showing higher tissue CX3CL1 signal in *Cx3cr1*^-/-^ mice compared to *Cx3cr1*^+/-^ mice. **e**. Graphical representation of the tissue CX3CL1 signal in *Cx3cr1*^+/-^ and *Cx3cr1*^-/-^ mice (n = 3 *Cx3cr1*^+/-^ mice and 4 *Cx3cr1*^-/-^ mice; Unpaired two-tailed t-test; Error bars represent SEM).

Additionally, we noticed a separate signal comprised of bright distinct puncta outside of the neuronal-associated labeling (Figure 1a; gray arrowheads). Given that microglia are the only known CNS cells to have a receptor that binds CX3CL1^9,10^ we hypothesized that these puncta represented CX3CL1 protein associating with the CX3CR1 receptor on microglia. Therefore, we co-stained coronal brain sections with an antibody against CX3CL1 and the microglia marker, IBA1. We observed that the bright discrete CX3CL1 puncta indeed associated with IBA1-positive microglial processes and somas, consistent with a ligand-receptor interaction under physiological conditions (Figure 1b and Supplementary Figure 2a-c). To determine if the CX3CL1 puncta on microglia were dependent on CX3CR1 being present on the microglia we performed immunostaining in *Cx3cr1*^-/-^ mice. The distinct puncta were indeed absent in the *Cx3cr1*^-/-^ tissues (Figure 1b-c and Supplementary Figure 2a-c). Moreover, in the absence of CX3CR1, CX3CL1 immunolabeling appeared markedly more diffuse with higher levels of extracellular signal (Figure 1b, d-e). Quantification confirmed a significant increase in CX3CL1 labeling within the tissue in *Cx3cr1*^-/-^ mice compared to the heterozygous controls (Figure 1d-e). This observation is consistent with previous reports showing that *Cx3cr1* deletion disrupts the primary clearance pathway for soluble CX3CL1, leading to its extracellular accumulation^33,34^.

### 2Phatal initiates neuronal cell death and microglial phagocytosis

To investigate the role of the CX3CL1/CX3CR1 signaling axis in microglial clearance of dying neurons in vivo, we used a previously established photochemical ablation technique for inducing cell death in single neurons, called 2Phatal^31,35–37^. Experiments were conducted in *Cx3cr1-creER*; floxed-tdTomato (Ai9) transgenic mice surgically implanted with dual cranial windows where the nuclei of all brain cells were labeled by applying nuclear dye (Hoechst 33342) directly to the brain pial surface during window implantation (Figure 2a). To induce neuron-specific cell death by 2Phatal, individual neuronal nuclei were selected based on morphological criteria consistent with neuronal identity i.e., relatively larger size and the presence of multiple nucleoli^31^. Selected neuronal nuclei were photobleached for ∼3 seconds to induce DNA damage and initiate cell death (Figure 2b, see methods). This photodamage resulted in nuclear pyknosis as observed by the condensation and bright fluorescence of the targeted cells by the 24-hour time point, a stereotyped characteristic feature of apoptotic progression in neurons with no evidence of cell rupture (Figure 2b). 2Phatal caused cell death in targeted neurons without damaging adjacent nuclei, illustrating the spatial precision of this method (Figure 2b). Selected neuronal nuclei were imaged before 2Phatal induced cell death, immediately after 2Phatal, then at 24- and 48-hours post 2Phatal. The efficiency rate of successfully inducing cell death across 683 targeted neurons was 54.47% ± 5.97% (Figure 2c).

**Figure 2:**
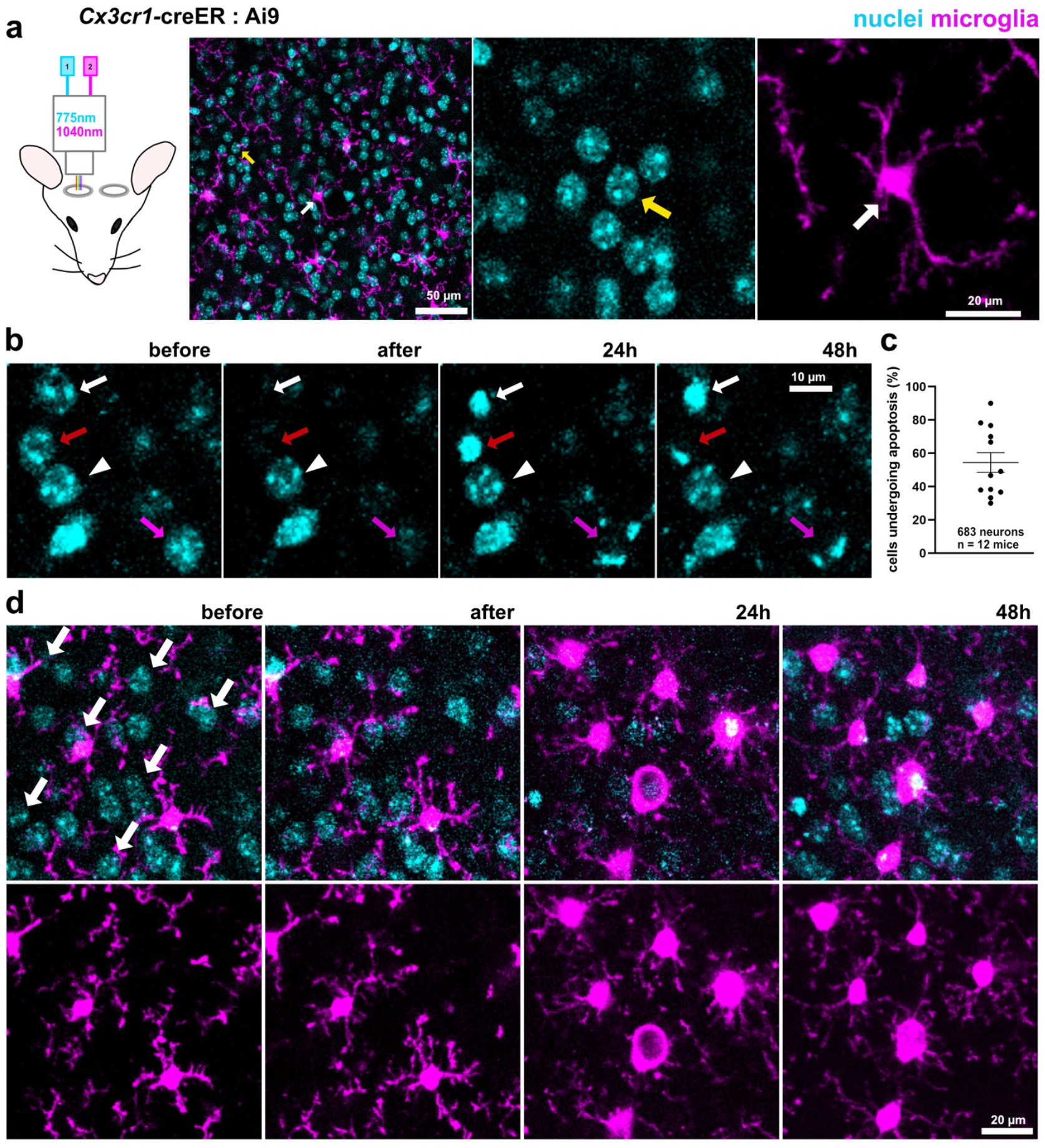
2Phatal-induced single-cell death in neurons leads to microglial engagement and clearance. **a**. Schematic diagram of 2-photon intravital imaging and representative images showing fluorescent labeling of all cell nuclei with Hoechst dye (yellow arrow) and microglia with TdTomato in the Ai9 reporter mice (white arrow). **b**. Representative examples of neuronal nuclei targeted with 2Phatal (all arrows) showing variability in the timing of cell clearance: cleared within 24 hours (magenta), cleared within 48 hours (red), and still present at 48 hours after 2Phatal (white). White arrowhead shows a neuronal nucleus that was not targeted with 2Phatal. **c**. Graph showing the 2Phatal success rate across all the targeted cells. Error bars represent SEM. **d**. Representative image showing the phenotypic shift of microglia from surveillant to phagocytic following successful induction of neuronal cell death with 2Phatal. The white arrows indicate targeted neuronal nuclei.

In positions where neurons were not targeted, microglia maintained their stereotyped highly ramified morphology and position in the parenchyma with no signs of chemotaxis (Supplementary Figure 3). However, following induction of 2Phatal in nearby neurons, microglia transitioned from their surveillant morphology to a phagocytic state coincident with nuclear condensation within the targeted neuron. This phenotypic shift was evident by 24 hours post-2Phatal, where microglia began to polarize and exhibit thicker shorter processes as they migrated toward the site of the dying neuron, features consistent with active engagement and engulfment of neuronal cell corpses (Figure 2d). These findings validate that 2Phatal not only initiated controlled, localized neuronal apoptosis but also reliably triggered a robust, spatially and temporally defined microglial response to dying neurons.

### CX3CR1 deletion impairs microglial engagement and clearance of single dying neurons

Our previous work implicated CX3CR1 as playing an important role in mediating microglial clearance of dying oligodendrocytes targeted by 2Phatal^29^. CX3CL1/CX3CR1 signaling has also been shown to mediate microglial clearance of dying neurons in a neonatal (P7) ethanol-induced apoptotic neuronal cell death model^28^. Whether this difference underscores a deficiency in the phagocytic capacity in the *Cx3cr1*^-/-^microglia or a broader change in the initial recruitment and engagement with the dying neurons is unclear. To study the role of CX3CR1 signaling axis in mediating microglial recruitment and clearance of dying neurons we used 2Phatal to target 415 neurons across 26 positions in 7 *Cx3cr1*^-/-^ mice and 268 neurons across 16 positions in 5 *Cx3cr1*^+/-^ control mice.

To analyze microglial behavior, we defined two distinct variables across all successfully targeted neurons in both genotypes. Previous studies characterizing the microglial clearance of dying neurons in vivo have shown that once apoptosis is initiated and find-me signals are released, microglia extend their processes toward the dying neuron forming an initial phagocytic cup before eventually migrating their somas to the region occupied by the dying neuron to fully phagocytose and clear the dying cell^36,38^. Thus, the first variable, which we termed “microglial engagement”, was the ability of microglia to find, chemotax, and contact the dying neuron (Figure 3a; white and orange arrowheads). The second variable, which we termed “microglial clearance”, was the successful phagocytosis and complete removal of the apoptotic neuron by microglia and is observed by the absence of nuclear dye (Figure 3a; white arrowheads). We restricted our analysis to the neurons we targeted that subsequently underwent apoptosis across the two genotypes (Figure 3a white arrows, *Cx3cr1*^+/-^: 133 neurons; *Cx3cr1*^-/-^: 233 neurons) to look at how deleting the microglial receptor, CX3CR1, affect microglia’s “find-me” and “eat-me” abilities. Analysis of these variables revealed a significant impairment in both the microglial engagement and clearance in the *Cx3cr1*^-/-^ mice when compared to the control *Cx3cr1*^+/-^ mice (Figure 3b). Importantly, the difference was not a secondary effect of unequal 2Phatal targeting efficiency across the two genotypes (52.25% ± 9.06% in *Cx3cr1*^+/-^ mice and 56.06% ± 8.47% in *Cx3cr1*^-/-^ mice; Figure 3c).

**Figure 3:**
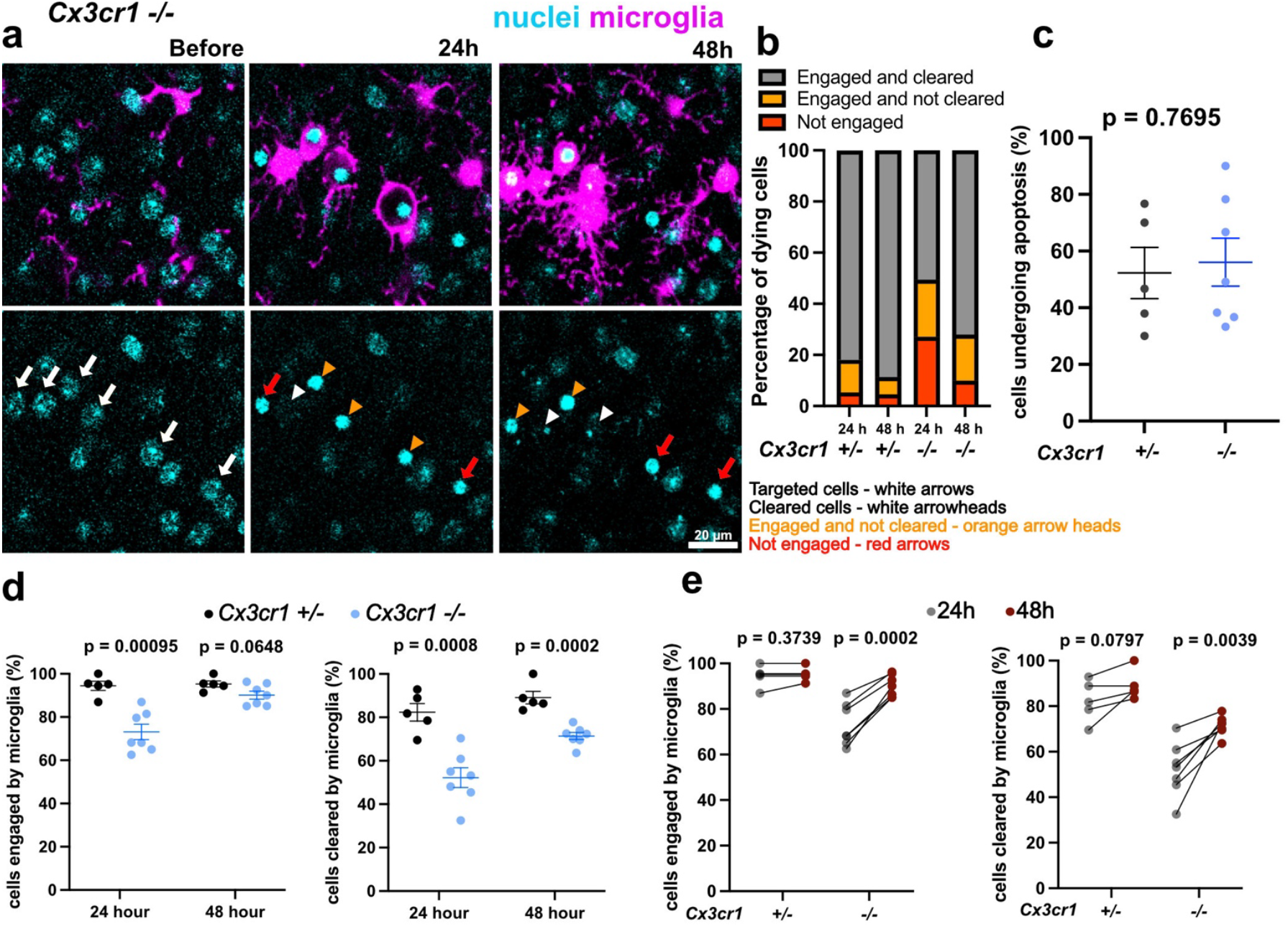
Microglial engagement and clearance of dying neurons is delayed in *Cx3cr1*^**-/-**^ microglia. **a**. In vivo images of microglial engagement (orange and white arrow heads) and microglial clearance of dying neurons following targeting by 2Phatal. The white arrows show targeted neuronal nuclei. **b**. Graph showing higher efficiency of microglial engagement and clearance in *Cx3cr1*^+/-^ microglia compared to *Cx3cr1*^-/-^ microglia at the 24- and 48-hour timepoints. **c**. The 2Phatal success rate in *Cx3cr1*^+/-^ and *Cx3cr1*^-/-^ microglia (n = 5 *Cx3cr1*^+/-^ mice and 7 *Cx3cr1*^-/-^ mice; Unpaired two-tailed t-test; Error bars represent SEM). **d**. The total cells engaged (left) and cleared (right) by microglia expressed as a percentage in Cx*3cr1*^+/-^ versus *Cx3cr1*^-/-^ mice at the 24- and 48-hour timepoints (n = 5 *Cx3cr1*^+/-^ mice and 7 *Cx3cr1*^-/-^ mice; Unpaired two-tailed t-test; Error bars represent SEM). **e**. The total cells engaged (left) and cleared (right) by microglia expressed as a percentage in *Cx3cr1*^+/-^ and *Cx3cr1*^-/-^ mice at the 24-vs 48-hour timepoints (n = 5 *Cx3cr1*^+/-^ mice and 7 *Cx3cr1*^-/-^ mice; Paired two-tailed t-test).

At the 24-hour timepoint, both the microglial engagement and clearance of dying neurons were significantly impaired in the *Cx3cr1*^-/-^ mice compared to the *Cx3cr1*^+/-^ controls (engagement: 73.12% ± 3.56% vs 94.42% ± 2.10%; clearance: 52.22% ± 4.54% vs 82.34% ± 4.07%; Figure 3d). By the 48-hour mark, however, the deficit in microglial engagement had resolved, with no significant difference observed between the genotypes (90.14% ± 1.83% in *Cx3cr1*^-/-^ mice vs 95.29% ± 1.39% in *Cx3cr1*^+/-^ mice; Figure 3d). In contrast, microglial clearance remained significantly impaired in the *Cx3cr1*^-/-^ mice even at 48 hours (71.41% ± 1.66% vs 89.11% ± 2.87%; Figure 3d).

Comparing across timepoints, we observed that both the microglial engagement and clearance of dying neurons increased significantly in the *Cx3cr1*^-/-^ mice between the 24- and 48-hour timepoints, whereas neither measure changed significantly across timepoints in the *Cx3cr1*^+/-^controls (Figure 3e). Altogether, these results show that knocking out *Cx3cr1* resulted in a significant delay in both the microglial detection and microglial clearance of dying cortical neurons in vivo.

### CX3CR1 deletion delays microglial clearance of dying neurons at multiple scales of cell death

To further explore the role of CX3CR1 in mediating microglial recruitment and clearance, we sought to investigate how disrupting this signaling axis would influence microglial functioning at different scales of cell death. A key limitation in the methods used to induce cell death in current literature is the inability to control the extent of cell death with previous studies often focusing on conditions with widespread cell death. These conditions might miss early stages of disease where widespread neurodegeneration is yet to occur and where therapeutics would be most beneficial. Additionally, a recent study using 2Phatal showed that increasing cell death led to a marked reduction in corpse removal efficiency, suggesting that microglial clearance efficiency may be sensitive to the overall burden of cell death (Rai et al., 2025). Thus, to take advantage of the spatiotemporal control available with the 2Phatal approach and to investigate the role of CX3CR1 signaling across different loads of cell death, we designed two distinct scales of neuronal cell death: a large-scale cell death model where we targeted 25 neurons and a small-scale cell death model where we targeted 5 neurons (Supplementary Figure 4 and 5).

In the large-scale cell death model, we targeted 228 neurons across 9 positions in 5 *Cx3cr1*^+/-^ control mice and 350 neurons across 14 positions in 7 *Cx3cr1*^-/-^ mice. Previous studies using different models of widespread neuronal cell death reported that deletion of the CX3CR1 receptor or its respective ligand led an increase in neuronal cell corpses^28^. Similarly, we observed that both the microglial engagement and clearance were significantly impaired in the *Cx3cr1*^-/-^mice (Figure 4a-b). Again, the observed difference was not due to a higher targeting efficiency in the *Cx3cr1*^-/-^mice (52.32% ± 9.02% for *Cx3cr1*^+/-^ and 58.86% ± 10.45% for *Cx3cr1*^-/-^; Figure 4c). At the 24-hour timepoint, both microglial engagement and clearance at this scale of cell death were significantly impaired in the *Cx3cr1*^-/-^ mice compared to the *Cx3cr1*^+/-^ controls (engagement: 74.08% ± 4.53% vs 93.20% ± 2.74%; clearance: 53.87% ± 5.20% vs 81.57% ± 4.93; Figure 4d). However, by the 48-hour timepoint, the difference in engagement was resolved (91.17% ± 4.53% vs 94.32% ± 2.74%; Figure 4d) while the difference in clearance persisted (73.34% ± 1.61% vs 88.60% ± 3.47; Figure 4d). Moreover, comparing across timepoints revealed that both microglial engagement and clearance significantly increased between the 24- and 48-hour timepoints in the *Cx3cr1*^-/-^ mice while neither measure changed significantly in the *Cx3cr1*^+/-^ controls (Figure 4e).

**Figure 4:**
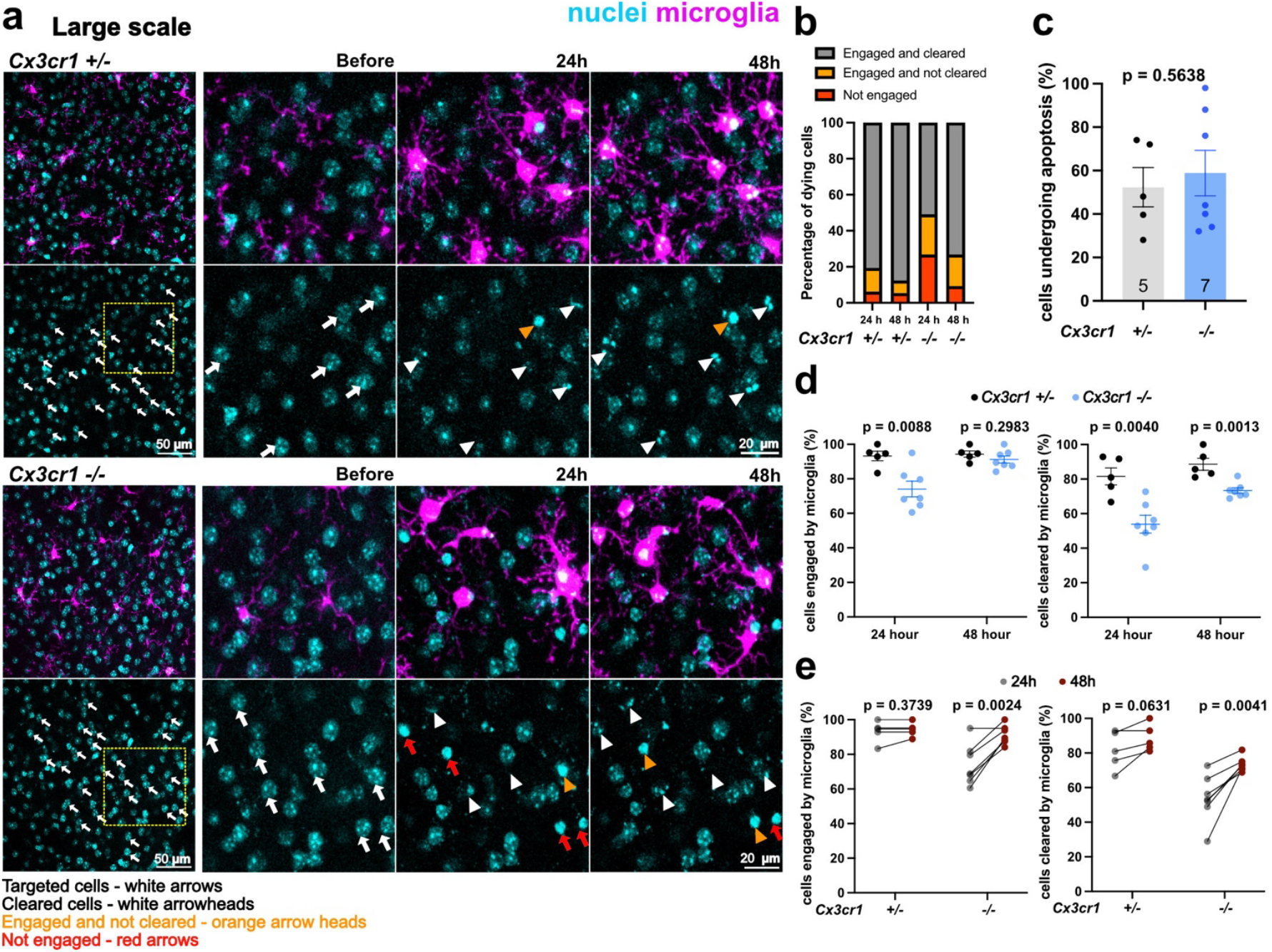
Microglial engagement and clearance of many dying neurons is delayed in *Cx3cr1*^**-/-**^ mice. **a**. In vivo images showing microglial engagement and clearance of dying neurons in the *Cx3cr1*^+/-^ controls compared to the *Cx3cr1*^-/-^ mice. **b**. Graph showing higher efficiency of microglial engagement and clearance in *Cx3cr1*^+/-^ microglia compared to *Cx3cr1*^-/-^ microglia at the 24- and 48-hour timepoints. **c**. Graph of the 2Phatal success rate in *Cx3cr1*^+/-^ and *Cx3cr1*^-/-^microglia (n = 5 *Cx3cr1*^+/-^ mice and 7 *Cx3cr1*^-/-^ mice; Unpaired two-tailed t-test; Error bars represent SEM). **d**. Graphs of the total cells engaged (left) and cleared (right) by microglia expressed as a percentage in *Cx3cr1*^+/-^ versus *Cx3cr1*^-/-^ mice at the 24- and 48-hour timepoints (n = 5 *Cx3cr1*^+/-^ mice and 7 *Cx3cr1*^-/-^ mice; Unpaired two-tailed t-test; Error bars represent SEM). **e**. Graphs of the total cells engaged (left) and cleared (right) by microglia expressed as a percentage in *Cx3cr1*^+/-^ and *Cx3cr1*^-/-^ mice at the 24-vs 48-hour timepoints (n = 5 *Cx3cr1*^+/-^ mice and 7 *Cx3cr1*^-/-^ mice; Paired two-tailed t-test).

We next performed the same experiments and analysis except we targeted 5 neurons instead of 25 in the condition termed small scale. Consistent with the large-scale conditions, the microglial engagement and clearance were impaired in the *Cx3cr1*^-/-^ mice compared to the *Cx3cr1*^+/-^ control mice (Figure 5a-e). Again, the observed difference was not due to a higher targeting efficiency in the *Cx3cr1*^-/-^ mice (50.00% ± 13.42 in *Cx3cr1*^+/-^ mice and 45.71% ± 10.43 in *Cx3cr1*^-/-^ mice; Figure 5c).

**Figure 5:**
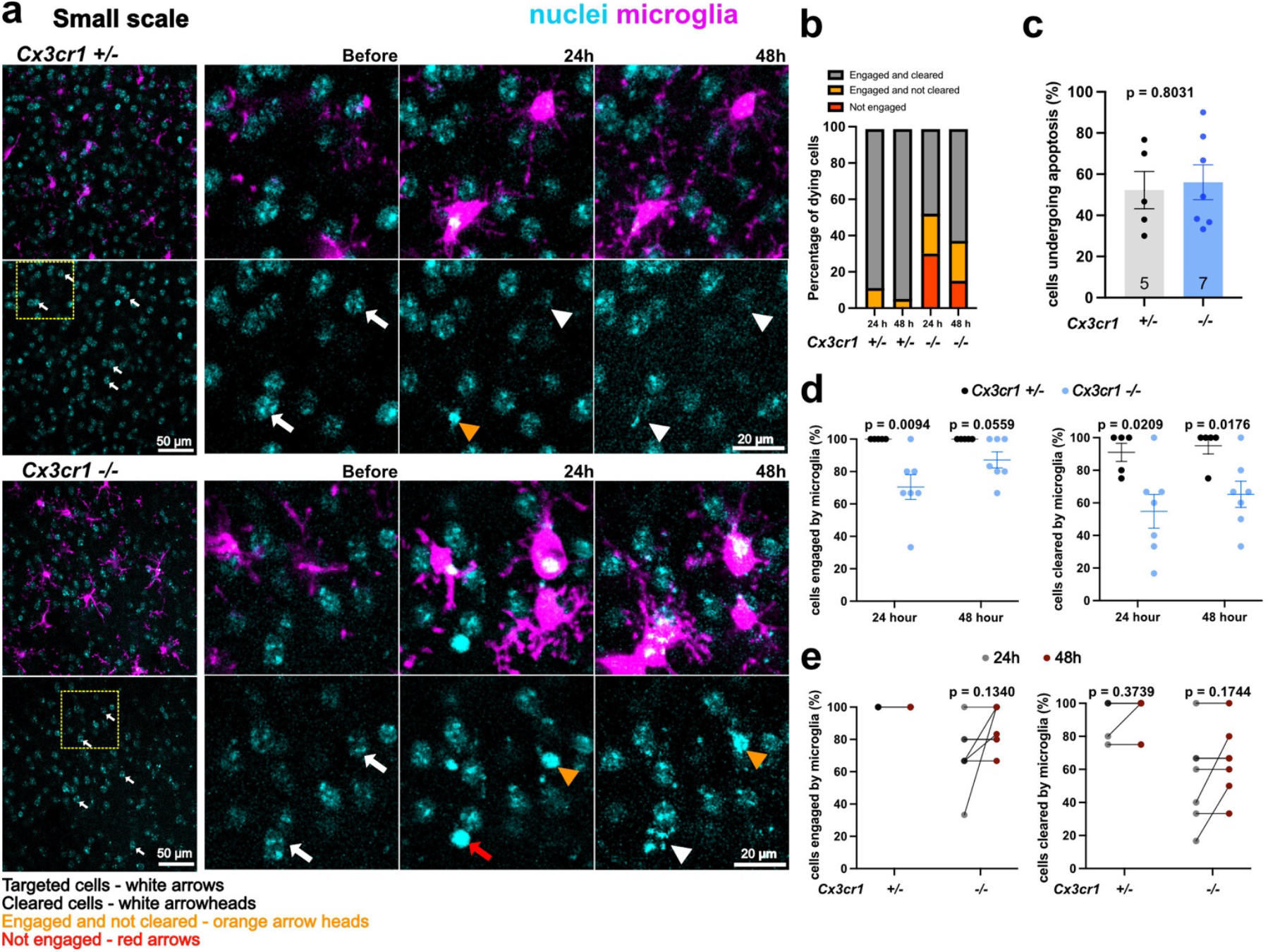
Microglial engagement and clearance of few dying neurons is delayed in *Cx3cr1*^**-/-**^ mice. **a**. In vivo images showing microglial engagement and clearance of dying neurons in the *Cx3cr1*^+/-^ controls compared to *Cx3cr1*^-/-^ mice. **b**. Graph showing higher efficiency of microglial engagement and clearance in *Cx3cr1*^+/-^ microglia compared to *Cx3cr1*^*-/-*^ microglia at the 24- and 48-hour timepoints. **c**. Graph of the 2Phatal success rate in *Cx3cr1*^+/-^ and *Cx3cr1*^-/-^ microglia (n = 5 *Cx3cr1*^+/-^ mice and 7 *Cx3cr1*^-/-^ mice; Unpaired two-tailed t-test; Error bars represent SEM). **d**. Graphs of the total cells engaged (left) and cleared (right) by microglia expressed as a percentage in *Cx3cr1*^+/-^ versus *Cx3cr1*^-/-^ mice at the 24- and 48-hour timepoints (n = 5 *Cx3cr1*^+/-^ mice and 7 *Cx3cr1*^-/-^ mice; Unpaired two-tailed t-test; Error bars represent SEM). **e**. Graphs of the total cells engaged (left) and cleared (right) by microglia expressed as a percentage in *Cx3cr1*^+/-^ and *Cx3cr1*^-/-^ mice at the 24-vs 48-hour timepoints (n = 5 *Cx3cr1*^+/-^ mice and 7 *Cx3cr1*^-/-^ mice; Paired two-tailed t-test).

Given that our 2Phatal targeting efficiency was not absolute, we were able to naturally obtain a spectrum of neuronal cell death events ranging from the death of just a single cell to as many as 25 across multiple fields of view. This variability allowed us to examine how both microglial engagement and clearance scaled with increasing levels of neuronal cell death. At the 24-hour timepoint, we observed that microglial engagement and clearance increased proportionally with the extent of cell death in both the *Cx3cr1*^+/-^ and *Cx3cr1*^-/-^ mice (Figure 6a-b). However, the rate of increase in both microglial engagement and clearance in the *Cx3cr1*^-/-^mice was consistently slower than the *Cx3cr1*^+/-^ mice (Fig. 6c).

**Figure 6:**
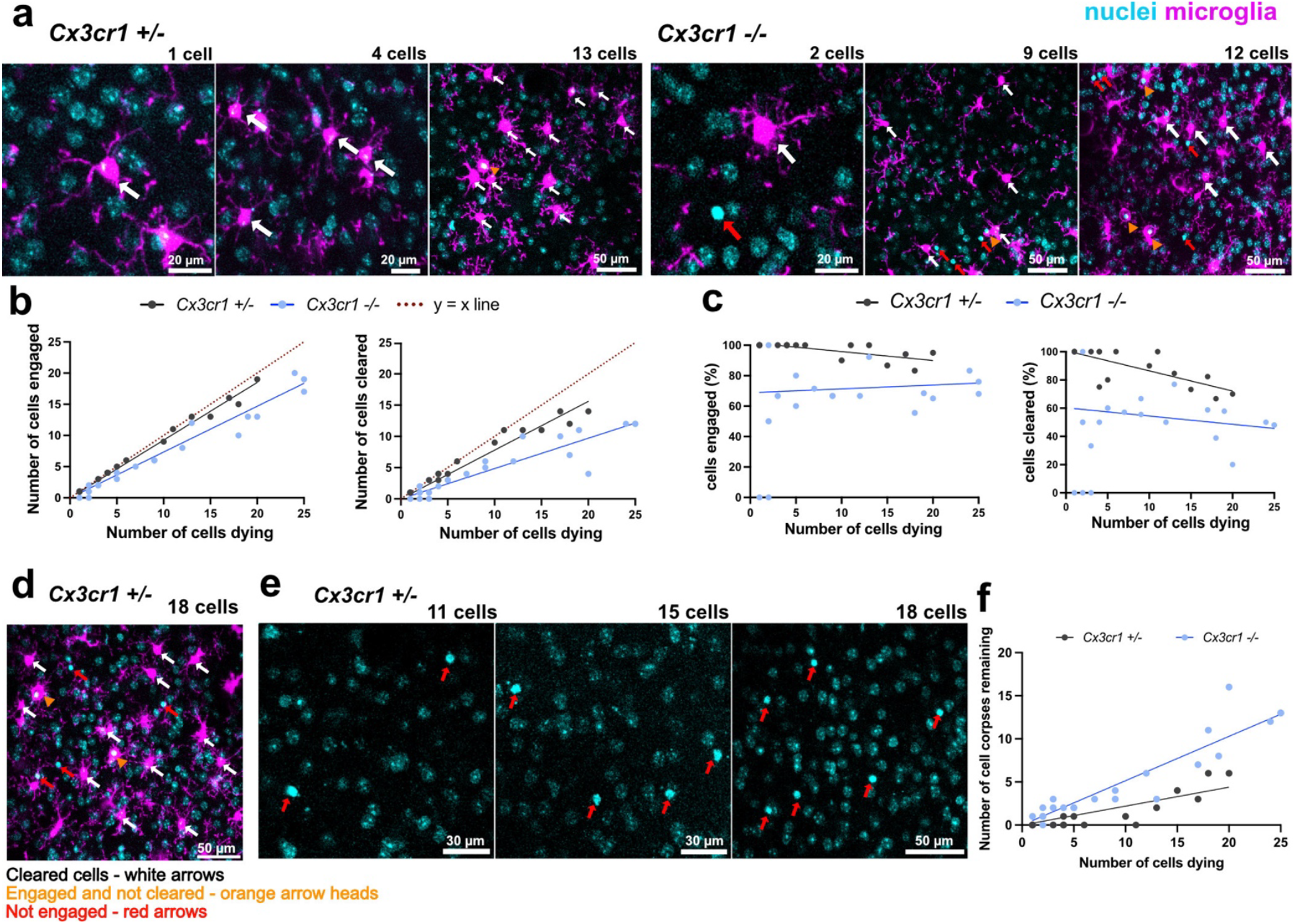
Titrating the burden of neuronal cell death reveals slower microglial dynamics in Cx3cr1^**-/-**^ mice. **a**. In vivo images showing microglial engagement and clearance scale slower in *Cx3cr1*^-/-^ mice (right panels) than in *Cx3cr1*^+/-^mice (left panels). **b**. Scaling of microglial engagement and clearance in *Cx3cr1*^-/-^ and *Cx3cr1*^+/-^ mice (n = 26 positions in 7 *Cx3cr1*^-/-^ mice and n = 16 positions in 5 *Cx3cr1*^+/-^ control mice). **c**. Graphs showing both microglial engagement and clearance are less efficient in *Cx3cr1*^-/-^ mice than in *Cx3cr1*^+/-^ mice (n = 26 positions in 7 *Cx3cr1*^-/-^ mice and n = 16 positions in 5 *Cx3cr1*^+/-^control mice). **d**. Representative image showing at high neuronal cell death burden, microglia clearance and engagement is less efficient. **e**. Representative images showing the number of remaining neuronal cell corpses increases with the number of cells dying. Red arrows show neuronal cell corpses not cleared at the 24-hour time point. **f**. The number of remaining cell corpses increases with increasing cell death (n = 26 positions in 7 *Cx3cr1*^-/-^ mice and n = 16 positions in 5 *Cx3cr1*^+/-^ control mice).

Additionally, as we previously noted, at the 24-hour timepoint *Cx3cr1*^+/-^ microglia exhibited highly efficient engagement approaching a near 1:1 ratio of dying cells to engaged microglia (∼100%) when fewer than 10 cells were dying (Figure 6a,c left panels). However, as the number of dying cells increased, engagement efficiency declined, suggesting that microglia approach a saturation point under high cell corpse burden (Figure 6c left panel and Figure 6d). A similar trend was observed for clearance efficiency, which was initially high but declined with increasing corpse load in the *Cx3cr1*^+/-^ mice (Figure 6c right panel). This was reflected in the accumulation of uncleared cell corpses in the *Cx3cr1*^+/-^ mice at 24 hours where nearly all dying cells were cleared when fewer than 10 cells were dying (Figure 6f). However, when more neurons were dying, there were more corpses (Figure 6e-f).

In contrast, the *Cx3cr1*^-/-^ mice showed a sustained impairment in both microglial engagement and clearance when only a few cells, as were dying which was maintained with more cell death (Figure 6c). This was most evident by the trend in corpse accumulation, with high numbers of uncleared cells even at low levels of neuronal cell death which then progressively worsened with increasing cell death (Figure 6f). Together, these results further show that *Cx3cr1* deletion delays both microglial engagement and clearance of dying neurons in vivo.

### CX3CL1 is present on microglial processes engaging a dying neuron

Following these observations, we next analyzed CX3CL1 localization when microglia respond to neuronal cell death. We performed multifocal 2Phatal of cells across both hemispheres in a young adult *Cx3cr1*^+/-^mouse, targeting 375 cells (Figure 7a, see methods). We then perfused the mouse immediately after and co-stained for CX3CL1, IBA1, nuclei-dye, and the Nissl stain (Neurotrace). Since CX3CL1 cleavage occurs early in the cell death process^15^ we focused our analysis on microglial processes extending out toward the dying neuron forming the initial phagocytic cup. We observed that microglial processes expand significantly during the formation of the initial phagocytic cup compared to neighboring processes surveilling nearby healthy control cells that were not targeted by 2Phatal (Figure 7a-c). Additionally, the increase in microglial process surface area was accompanied by an increase in the number of fractalkine associated puncta (Figure 7d). When the CX3CL1 signal is normalized to the total microglial process signal there is no difference (Figure 7d). This shows that as microglial processes initiate phagocytosis, they undergo membrane remodeling to maintain their total surface CX3CR1 receptors. Taken together, these results show that soluble fractalkine is cleaved and released by neurons during both physiological and pathological conditions which binds its cognate receptor, CX3CR1, on microglia.

**Figure 7:**
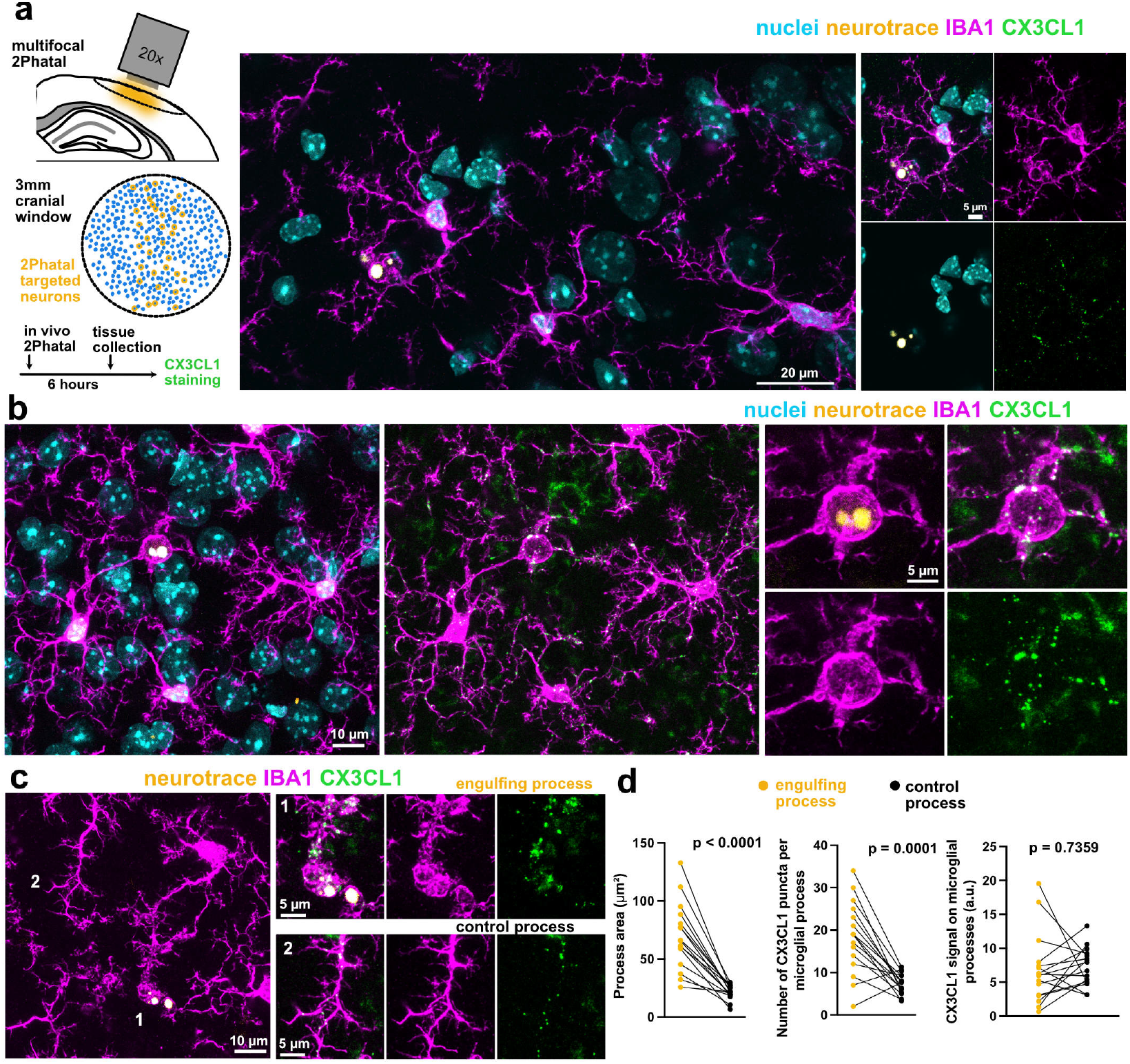
CX3CL1 binds CX3CR1 on microglial processes engaging dying cells. **a**. Left: Schematic diagram of multifocal 2Phatal targeting of neurons across the mouse neocortex and a timeline of experimental procedures. Right: Representative image of an IBA1^+^ microglia engulfing a dying cell labeled with Neurotrace **b**. Representative images showing an expanding an IBA1^+^ microglial process forming a phagocytic cup as it engulfs a dying cell labeled with Neurotrace. **c**. Representative images showing CX3CL1 puncta is maintained when microglia processes are in surveillance and when microglial processes are engulfing a dying cell. **d**. Greater process area and number of CX3CL1 puncta in engulfing processes compared to neighboring control processes. The increase in both features results in no difference in the relative CX3CL1 signal on the microglial processes (n = 16 processes; Paired two-tailed t-test).

## DISCUSSION

The CX3CL1/CX3CR1 signaling axis is increasingly recognized as a context-sensitive regulator of microglial function. This signaling ranges from homeostatic maintenance to both protective and detrimental outcomes in various pathological models depending on timing and disease progression^39^. Here, we applied a non-inflammatory single-cell apoptosis model in combination with longitudinal 2-photon in vivo imaging to determine how CX3CR1 mediates microglial responses to different extents of neuronal death.

We show that CX3CR1 is not necessary for the eventual microglial clearance of dying neurons in vivo but rather it modulates the rate of microglial responses to neuronal death. At multiple scales of cell death, *Cx3cr1* deletion led not only to a delay in the overall microglial clearance but also to a delay in microglial engagement with the dying neurons, particularly within the first 24 hours following targeting by 2Phatal. These findings suggest that the impaired clearance observed in the *Cx3cr1*^-/-^mice is not simply a failure in phagocytosis but rather, at least in part, is due to a delay in the microglial detection of the dying neurons. CX3CL1 is well established to be a chemoattractant for microglia mediating their migration via CX3CR1^9,15,40^. In the context of cell death, CX3CL1 cleavage and release has been shown to occur quickly after cell death induction and outpace the rate of apoptosis as seen in a study of dying Burkitt Lymphoma cells targeted by UV radiation and pharmacology^41^ as well as in a study of dying neuronal cells induced by glutamate excitotoxicity^15^. Cell culture supernatants from the neuron study were sufficient to induce migration of CX3CR1-expressing cells suggesting that CX3CL1 is cleaved by dying neurons early in the cell death process as a “find-me” signal. Our results are consistent with these findings by showing that CX3CL1 is indeed present on microglial processes both at baseline and when engaging a dying cell. More importantly, we found that at small scales of cell death, where the loss of a single chemotactic molecule would potentially have the greatest effect, microglial engagement with dying neurons is indeed significantly delayed in *Cx3cr1*^-/-^ mice. Given the range of other documented “find-me” signaling mechanisms present on neurons and microglia^42^, it is notable that the deletion of CX3CR1 alone was sufficient to have an effect on cell corpse clearance.

In addition to a direct role for CX3CL1 signaling from a dying neuron to local microglia, a CX3CR1-dependent defect in microglial process and migratory kinetics could also lead to the observed delay in microglial engagement and clearance. In the retina, *Cx3cr1* deletion results in slower microglial surveillance and responses to a focal laser lesion^43^. In the cortex, however, *Cx3cr1* deletion causes enhanced microglia migration and increased length and velocity of microglial processes extending toward a laser lesion^44^. Reasons for these contradictory data are not clear, however, retinal microglia have been shown to possess specialized transcriptomic profiles, appearing more inflammatory and exhibiting higher MHC class I expression as compared to cortical microglia^45^ which could explain differences in the context of *Cx3cr1* deletion. Similarly, it is possible that the deletion of *Cx3cr1* alters cortical microglial gene expression patterns making them more sensitive to inflammatory injuries such as laser lesions or advanced neurodegenerative disease states but not as sensitive to non-inflammatory programmed cell death. Indeed, RhoA signaling, which has been associated with spontaneous microglial activation and increased expression of pro-inflammatory activity^46^, was downregulated in *Cx3cr1*^-/-^ mice^44^. Similarly, *Cx3cr1*^-/-^microglia have an increased pro-inflammatory signature^21,47,48^, exhibit shorter and less ramified processes^48,49^, and show increased phagocytic activity^21–23^, all indicative of a change in microglial state. Altogether, these studies show that *Cx3cr1* deletion does indeed alter the microglial state both molecularly and functionally, making them more attune to inflammatory contexts and potentially inefficient in non-inflammatory contexts such as apoptosis.

Another factor that may be affected following *Cx3cr1* deletion could be the debris degradation process within microglia. While we do not anticipate that degradation was affected by *Cx3cr1* deletion, it is important to note that our study did not directly assess this, thus whether the deficits observed in clearance are fully attributed to impaired detection or instead reflect a cumulative disruption across both detection and engulfment and/or degradation, remains an open question.

In summary, we show that neurons express and release CX3CL1 that binds to its respective receptor, CX3CR1, on microglia when microglial processes are adjacent to healthy, and engulfing dying, neurons. We also show that *Cx3cr1* deficiency significantly delays both microglial engagement and clearance of single-targeted cortical neurons at multiple scales of cell death, and that increasing the cell death burden can further slow microglia function. Altogether, these data demonstrate the significance of CX3CR1 signaling in maintaining homeostasis under early neurodegenerative contexts and suggest that the disruption of this pathway may shift the efficiency and timing of microglial detection and clearance of dying cells.

## METHODS

### Animals

All the animal-related procedures performed in this study were approved by Dartmouth College’s Institutional Animal Care and Use Committee (IACUC). The study used two transgenic mouse lines: *Cnp-mEGFP*; *Cx3cr1-creER*; floxed-tdTomato Ai9 and *Cx3cr1-creER*; floxed-tdTomato Ai9, which were generated by crossing the following mouse strains purchased from The Jackson Laboratory: Cx3cr1-creER (strain no. 020940)^50^, *Cnp-mEGFP* (strain no. 026105)^51^, and floxed-tdTomato Ai9 (strain no. 007909)^52^. In total, 12 mice (6 males and 6 females), aged 3-5 months at the start of the experiments, were used. Among them, 8 mice of the *Cnp-mEGFP, Cx3cr1-creER*, floxed-tdTomato Ai9 strain had oligodendrocytes labeled by excitable *Cnp-mEGFP*, though the fluorophore was not excited nor visualized for this study. All mice were housed in a temperature- and humidity-controlled vivarium with a 12-hour light/dark cycle (lights off at 7pm) and had ad libitum access to food and water through the entire length of the study. The number of mice used per experiment is specified in the figure legends.

### Tamoxifen Injections

For the purposes of this study, microglia were visualized via a red fluorescent protein variant tdTomato which was conditionally expressed through cre-mediated recombination. For this reason, all the mice used in the study were either homozygous or heterozygous *Cx3cr1*^*CreER*^ knockin/knockout mice where the fusion protein CreER was expressed under the endogenous *Cx3cr1* promoter thus knocking out endogenous *Cx3cr1* expression in at least one allele. To induce Cre mediated recombination, tamoxifen was dissolved in corn oil for a final concentration of 20 mg/mL. All mice were then injected with 0.05 mL of tamoxifen intraperitoneally for two consecutive days after weaning.

### Surgical Procedures

Dual cranial window implantations were performed following established protocols^31,36^ to allow for all subsequent in vivo imaging. Briefly, mice were anesthetized with Ketamine (100mg/kg) and Xylazine (10mg/kg). Before the procedure, the head was shaved and sterilized. The skin was removed overlying the skull and skull and dura matter were then removed to expose a 3-4 mm region of the brain cortex directly above the somatosensory cortex, which was subsequently replaced with a glass window. During the surgery, the nuclear dye Hoechst 33342 (10mg/mL) (Fisher, Invitrogen) was diluted 1:150 in sterile PBS and applied topically to the pial surface for in vivo nuclear labeling. Following the procedure, mice were allowed to recover and received carprofen (5 mg/kg) both pre- and post-operatively for pain management.

### 2Phatal

To target cells for death in this study, we used a single cell ablation technique called 2Phatal which is an abbreviated term for 2-photon chemical apoptotic targeted ablation^31,36,37,53^. This technique allows for precise nuclear photobleaching by using a topical dye, Hoechst 33342, which is applied during cranial window implantation and binds the DNA to selectively label the nuclei of all brain cells. Single cell death by 2Phatal was performed by placing a single 15 by 15 pixels region of interest (ROI) over the nucleus of the neuron of interest and photobleaching the nuclear dye using a 775 nm wavelength laser on the 2-photon microscope for approximately 3 seconds (125 scans) which is enough to induce sufficient DNA damage to trigger apoptosis in the neuron.

Multifocal 2Phatal^54^ was done to allow the identification and analysis of 2Phatal targeted cells, and the microglia that engulf them, in fixed tissue. The 2Phatal procedure was the same as described above except many cells across the entire cranial window were targeted. The targeted cells were purposefully distributed in the anterior to posterior regions of the cranial window to generate a substantial number of coronal tissue slices that could be stained, imaged, and analyzed. Targeted and dying cells were identified through a combination of condensed nuclei and bright Neurotrace dye labeling.

### Immunohistochemistry

For the fixed tissue stains performed in this study, mice were transcardially perfused with a solution of 4% paraformaldehyde (PFA) in phosphate-buffered saline (PBS). Isolated brains were post-fixed overnight at 4 degrees Celsius in 4% PFA then transferred to 1x PBS and stored at 4 degrees Celsius until sectioning. Sectioning was performed using a vibratome to slice tissues coronally at a thickness of 75µm. Sections were then cryo-preserved at -20 degrees Celsius. For the immunohistology experiments shown in this study, slices from the primary somatosensory cortex were selected, washed 3 times with sterile 1x PBS, then heated at 98 degrees Celsius with pre-heated antigen retrieval solution (10mM Tris, 1mM EDTA, 0.05% Tween buffer in PBS, at pH 9) for two minutes. Sections were then incubated with the primary antibodies overnight at room temperature in blocking solution (1% bovine serum albumin (Sigma, cat. No. A9418-5G), 0.3% Triton X-100 in sterile 1x PBS). The following day slices were washed 3 times with sterile 1x PBS then incubated with the respective secondaries in a blocking solution for two hours on rotation after which the sections were washed 2 times with PBS then incubated in nuclear dye and Nissl (Neurotrace) stain for thirty minutes. After this staining process, the slices were then mounted on slides using 40µL of Prolong Diamond Antifade Mountant (Catalog No. P36970, Invitrogen).

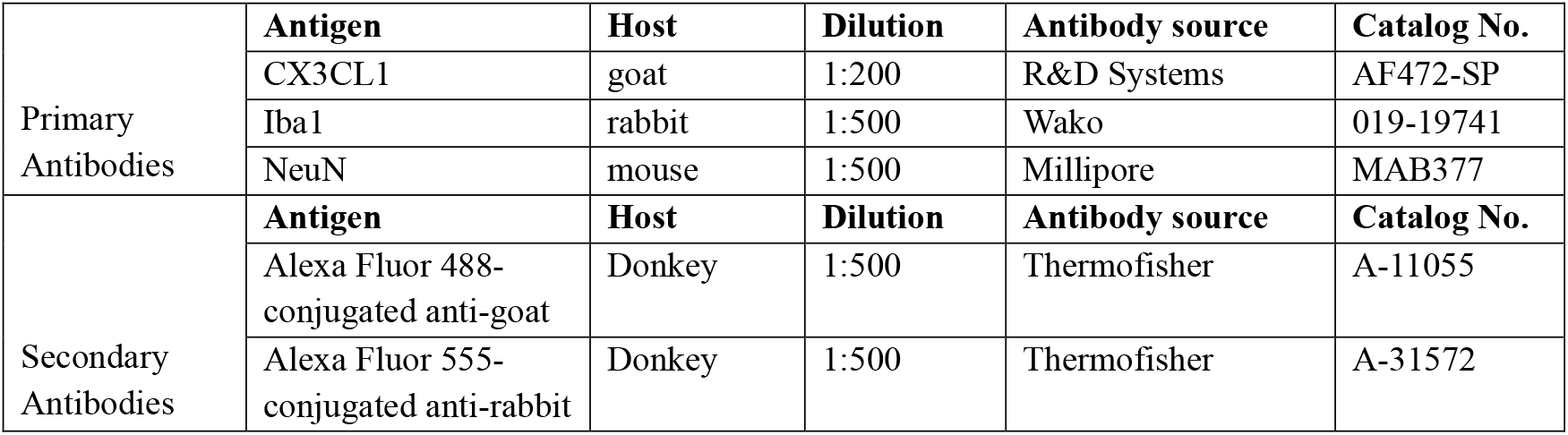

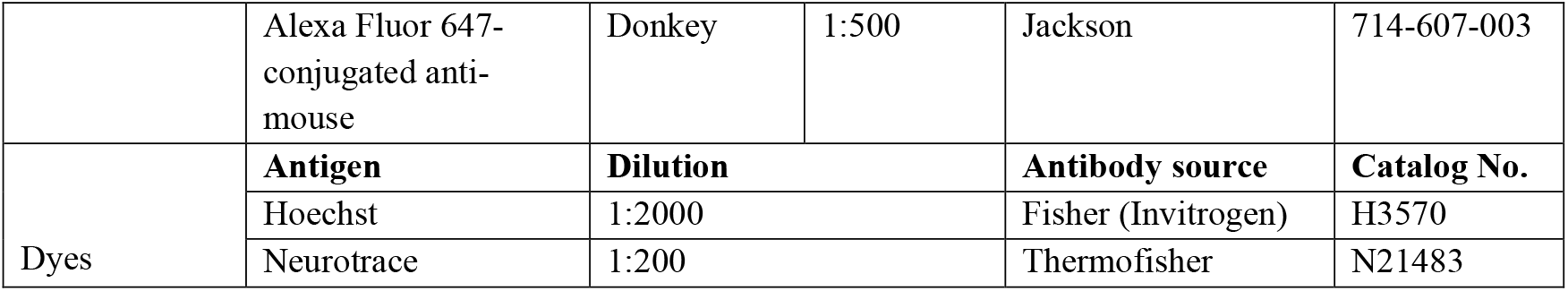

### Imaging

To visualize cells in vivo, mice were anesthetized using 3.0% isoflurane and anesthesia was maintained with 1.5% isoflurane using the SomnoSuite, Low-Flow Anesthesia System. Images were captured with the Prairie View software v.5.4 on a Bruker 2-photon microscope equipped with an Insight X3 femtosecond pulsed laser (Spectra Physics) and a 20x water immersion objective (Zeiss NA 1.0). Imaging was performed at depths of approximately 150 - 200 µm from the cortical surface, corresponding to layer II of the somatosensory cortex. The Hoechst dye was excited with the 775nm laser, and the tdTomato fluorophore was excited with the 1040nm laser. Z-stacks were captured at a 2x zoom, with a step size of 1.5 µm and at a 512 x 512 pixel resolution (272.97 x 272.97µm). For longitudinal imaging, blood vessels close to the surface were captured on day 0 in each region of interest to be used as landmarks for repeated imaging of the same position.

To visualize immunolabeled fixed tissue samples, imaging was conducted using the LasX software v.3.5.7 on an upright laser scanning confocal microscope (Leica SP8) with a 20x air objective (Leica NA 0.75) and a 63x oil immersion objective (Leica NA 1.4). Images were acquired sequentially using the 405nm, 488nm, 555nm, and 647nm lasers.

### In vivo image processing and quantification

In vivo brain images were analyzed using FIJI/Image J software (version: 2.16.0/1.54r). To quantify the efficiency of 2Phatal, the number of cell nuclei condensed or absent at the 24-hour timepoint were manually counted and compared to the total number of cells targeted with 2Phatal per position.

To quantify the percentage of cells engaged and/or cleared by microglia at the different timepoints, cells were counted manually according to the rationale explained in the results section and compared to the total number of cells that were targeted and condensed/ absent by the 24-hour timepoint per position.

### Fixed tissue image processing and quantification

Fixed tissue brain images were analyzed using FIJI/Image J software. To quantify the CX3CL1 signal on microglial processes, composite images were first duplicated and blinded prior to analysis. All fluorescent channels except the microglia fluorescent channel were temporarily turned off so that microglia could be visualized in isolation. Using an unbiased number generator, a microglial cell was randomly selected for analysis. From this microglial cell of interest, two processes were chosen, and the two z-planes in which fluorescence of each selected process was highest were used to generate a maximum intensity projection. A 20-µm wide line ROI was then used to trace the selected microglial process. After the ROI was defined, the mean gray value fluorescence of the CX3CL1 channel within the traced ROI was measured to obtain the raw process-associated CX3CL1 signal. To normalize for background signal, the same ROI was subsequently moved to three nearby regions of tissue that were devoid of nuclei and microglial processes and the mean fractalkine fluorescence in these regions was measured. The final process-associated CX3CL1 fluorescence was calculated by subtracting the average tissue-associated fluorescence from the raw process-associated fluorescence.

To quantify microglial process area, 20×20 µm cropped images were first generated and centered on the targeted condensed neuronal nuclei. All fluorescent channels except the microglia channel were temporarily turned off so that microglia could be visualized in isolation. The number of z steps required to fully capture microglial process engaging the dying neuron was determined, and only these slices were used to generate a restricted z-stack. Before generating a max projection, slices were examined individually with the CX3CL1 fluorescent channel temporarily turned on. Any CX3CL1 signal originating from nearby healthy neurons and not associated with the process of interest that could potentially overlap with the process of interest in other z-planes was manually removed using the selection brush tool. A maximum intensity projection was then generated from the edited stack. The channels were then split, and only the microglia and CX3CL1 channels were saved separately. The microglial channel was auto thresholded using “Otsu” to generate a binary mask of the process of interest, and the “Analyze Particles” function was used to generate an ROI corresponding to this process. In cases where the process formed a phagocytic cup, internal holes within the mask were manually traced using the selection brush tool to generate a second ROI. The “XOR” function was then applied to create a final ROI that excluded these holes from any subsequent measurements. The area of the microglial process of interest was then quantified using this ROI. Three control processes were randomly selected from neighboring microglia per position if the entire process was visible within the predetermined z-stack and the average microglial process area of the three was recorded for that position. All other analysis steps were maintained for the control processes, except that cropping was centered on a healthy neuronal nucleus.

To quantify the number of process-associated CX3CL1 puncta and the process-associated CX3CL1 signal, the previously saved CX3CL1 channel was opened in FIJI and auto thresholded using “Max Entropy”. The process-associated ROI generated from the microglial mask was then applied to the thresholded CX3CL1 image, and any signal outside this ROI was cleared to restrict the analysis to the microglial process. The “Analyze Particles” function was then used to generate individual ROIs corresponding to CX3CL1 puncta, with the particle size set between 0.05-Infinity (µm^2^). The total number of puncta within the process was then recorded. To quantify overall CX3CL1 signal associated with the process, the same process-ROI was applied to the CX3CL1 channel and the mean gray value fluorescence was measured.

### Statistical analysis

All statistical analyses were conducted using GraphPad Prism software version 10.3.0 (461). Multiple unpaired t tests and paired t tests were employed to determine significant differences between the different in vivo and fixed tissue conditions outlined in this study. Differences were considered statistically significant when the probability, p, of the null hypothesis was <0.05. Data are represented as mean +/-S.E.M. Final images were then prepared and organized using Affinity designer.

## ACKNOWLEDGEMENTS

We thank members of the Hill Lab for helpful discussions and feedback. This study was supported by grants from the National Institutes of Health R01NS122800 and R01NS140248 to R.A.H. and the E.E. Just Fellowship and the Thomas B. Roos Memorial Fund Fellowship to M.N.B.

## AUTHOR CONTRIBUTIONS

M.N.B. designed and performed all in vivo imaging experiments, immunostaining and fixed tissue imaging, analyzed the data, and wrote the first draft of the manuscript. A.N.P. contributed guidance on data analysis and editing of the manuscript. R.A.H. designed experiments, performed surgeries, provided guidance on data analysis, edited the manuscript, secured funding, and supervised the study.

