## Supplementary Figures for "Microglia detection and phagocytosis of dying neurons is regulated by CX3CR1"

### SUPPLEMENTARY INFORMATION

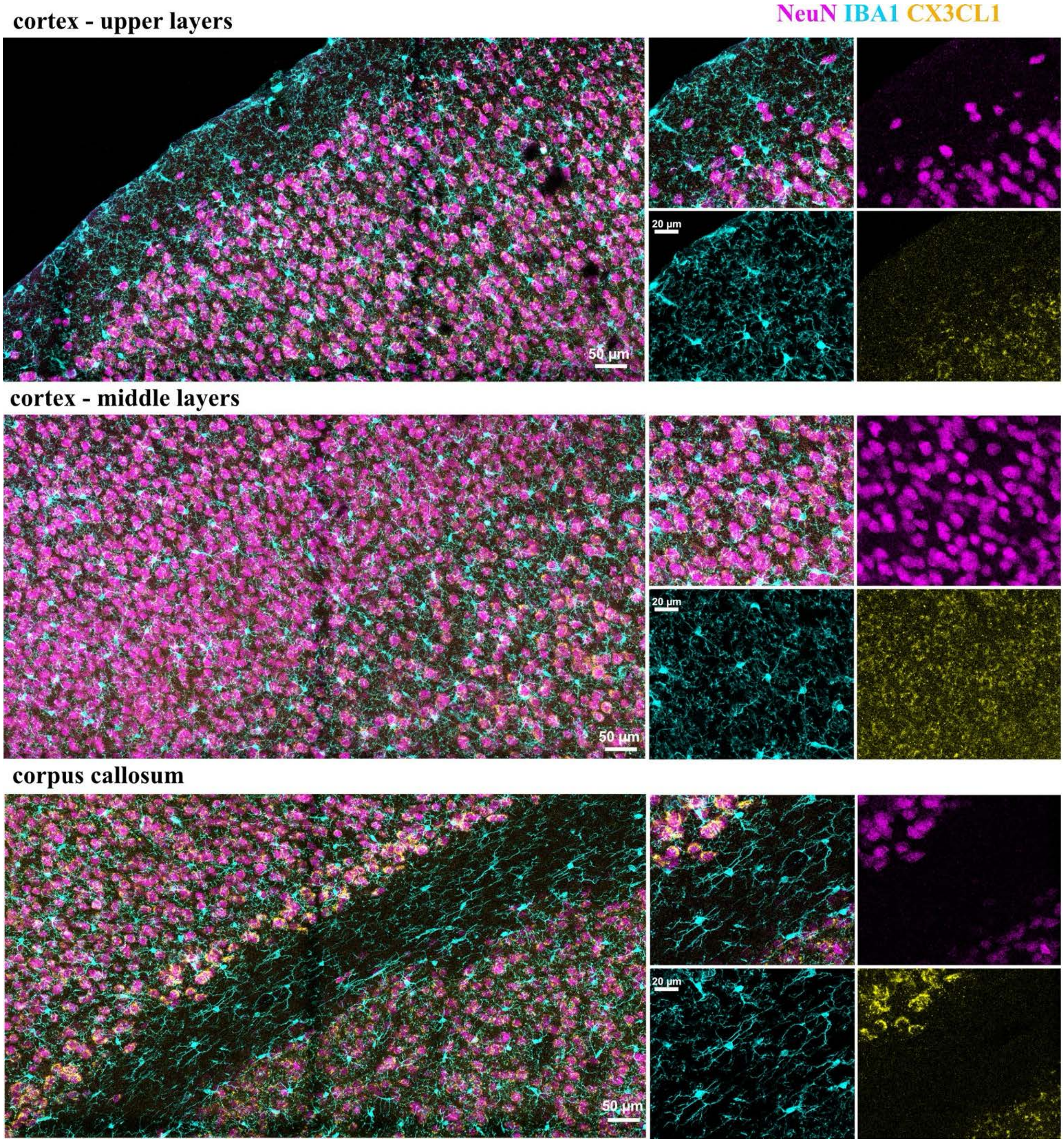


**Supplementary Figure 1: CX3CL1 localization in the mouse neocortex and corpus callosum.** Images showing CX3CL1 immunoreactivity (yellow) in the different layers of the mouse cortex and corpus callosum. CX3CL1 immunoreactivity is present in cortical layers II-VI colocalizing with $NeuN^+$ cells (magenta) and absent in cortical layer I and the corpus callosum where there is little to no NeuN immunoreactivity.

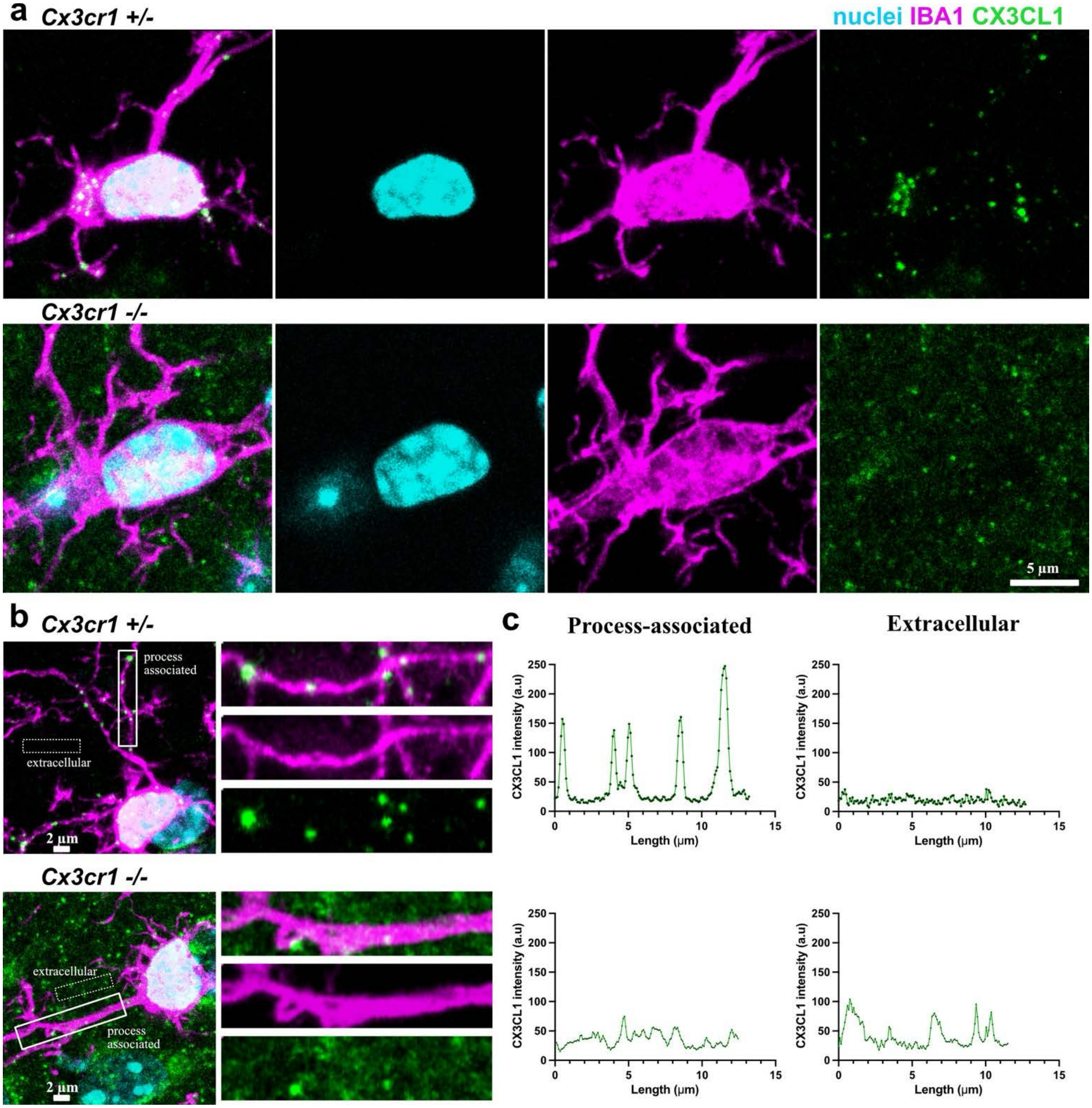


**Supplementary Figure 2: CX3CL1 puncta aggregation on microglia is dependent on CX3CR1. a.** CX3CL1 immunoreactivity colocalizes with IBA1 staining on the microglial cellular membrane as bright distinct puncta (top panel) which are lost when the CX3CR1 receptor is deleted (bottom panel). **b.** Example images of CX3CL1 fluorescent signal on microglial processes and in the extracellular matrix in the presence (top panels) and absence (bottom panels) of the CX3CR1 receptor. **c.** Graphical representation of the CX3CL1 fluorescent intensity in the boxed regions of the *Cx3cr1*$^{+/-}$ and *Cx3cr1*$^{-/-}$ microglia shown in (b).

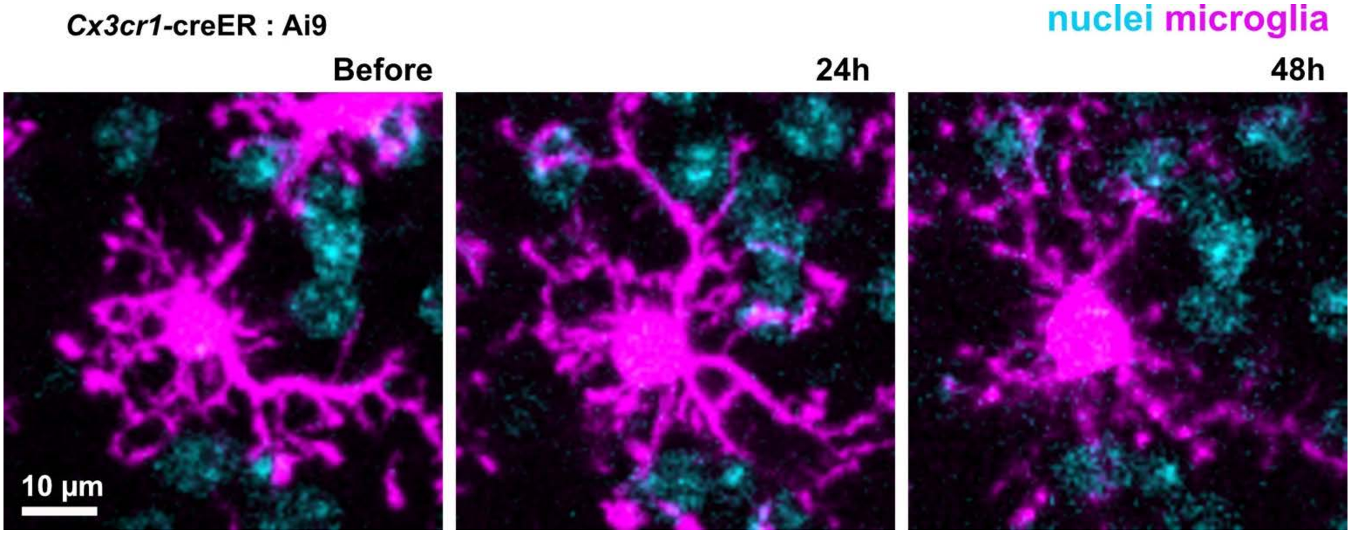


**Supplementary Figure 3. Surveillant microglia maintain a stationary soma while their processes remain highly dynamic in the absence of neurodegeneration.** Representative image showing a microglial cell imaged over a period of 48 hours away from any targeted cells labeled with TdTomato in the Ai9 reporter mice. The cell soma maintains its specific position in the parenchyma while its processes are seen to change at each timepoint, a characteristic of a surveillant phenotype.

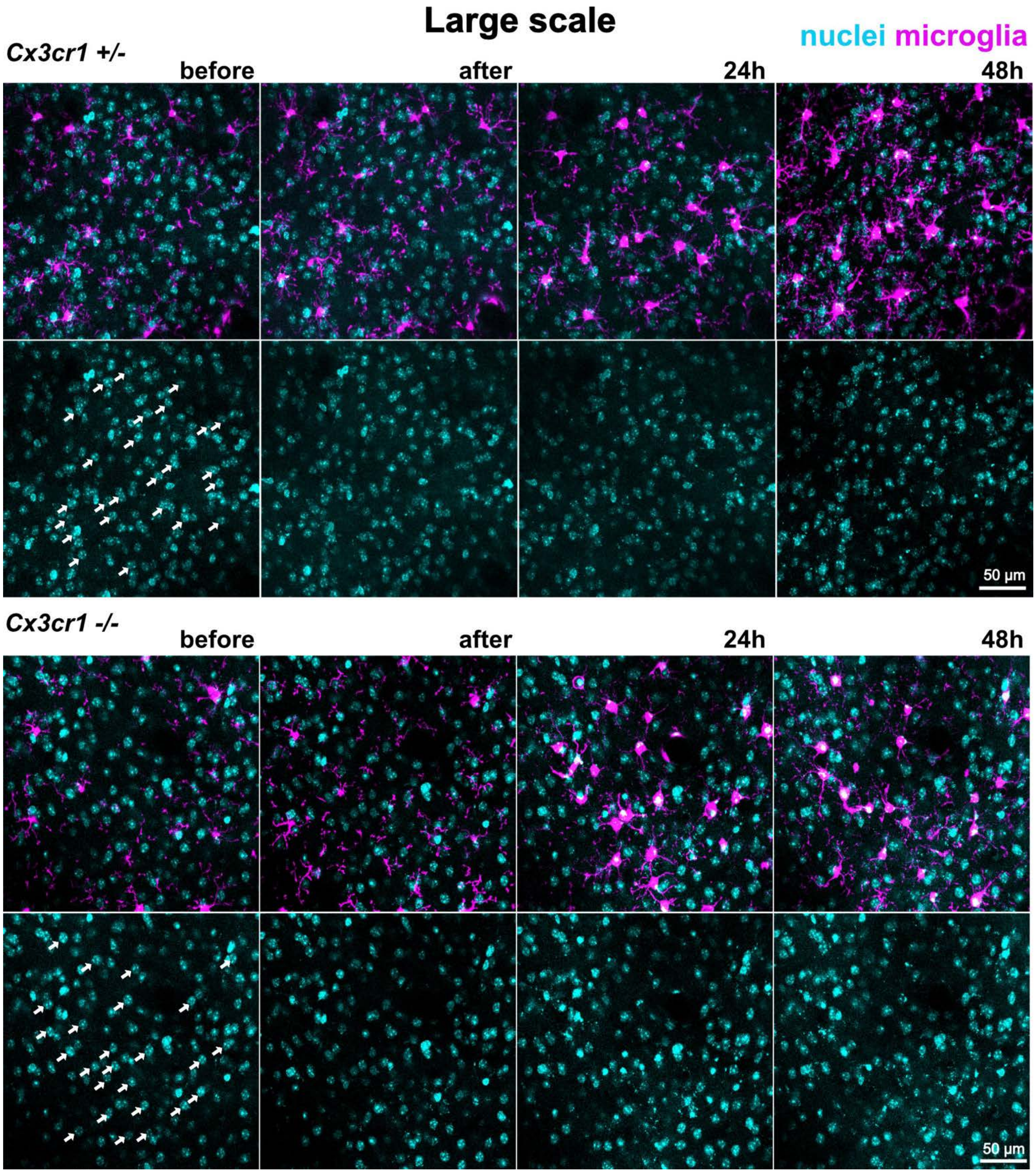


**Supplementary Figure 4. Representative image of longitudinal imaging of the large-scale cell death model.** Representative images showing twenty-five cells targeted per position in *Cx3cr1*$^{+/-}$ mice (top panels) and *Cx3cr1*$^{-/-}$ mice (bottom panels) across different timepoints. White arrows show targeted cells.

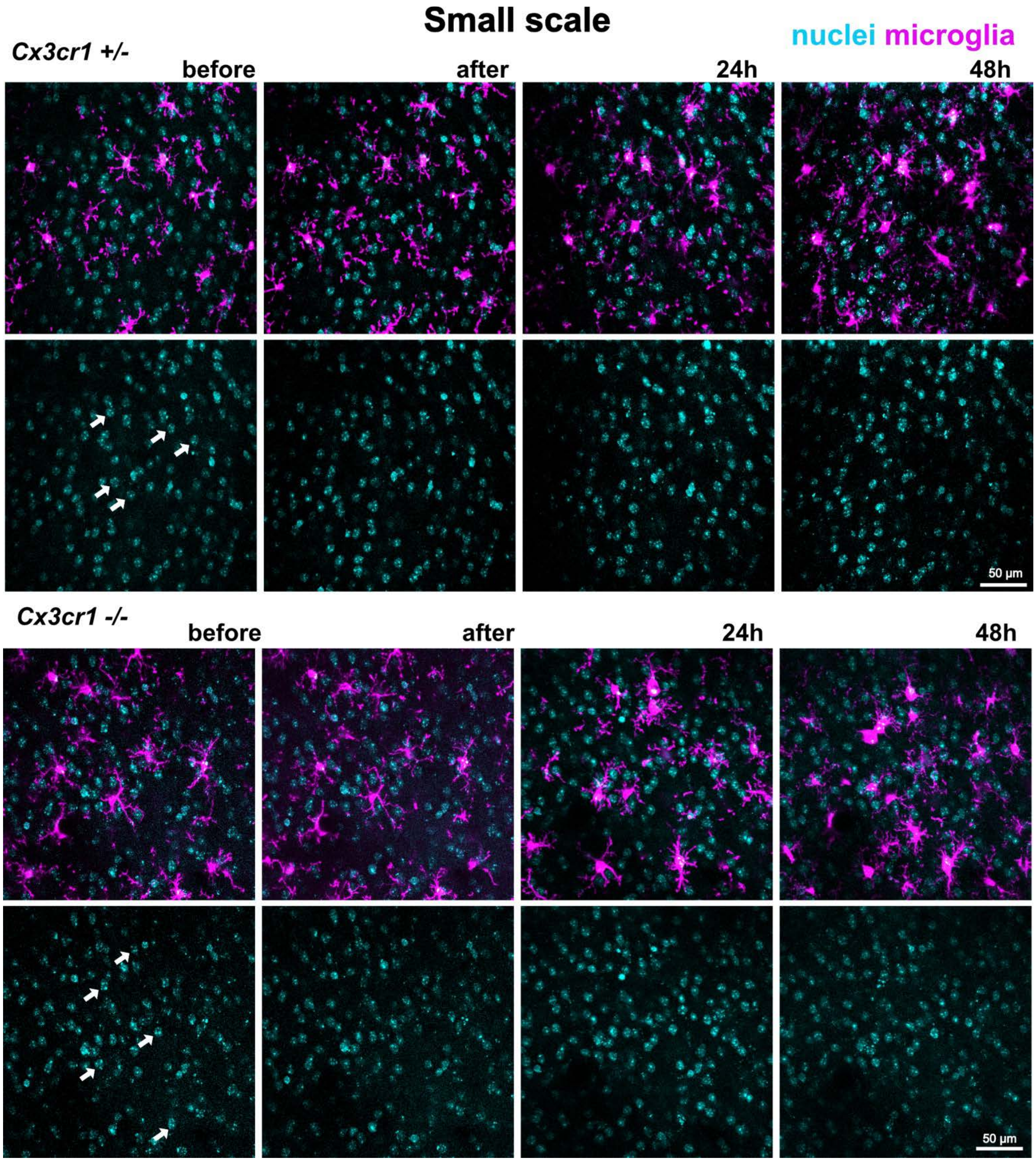


**Supplementary Figure 5. Representative image of longitudinal imaging of the small-scale cell death model.** Representative images showing five cells targeted per position in *Cx3cr1*$^{+/-}$ mice (top panels) and *Cx3cr1*$^{-/-}$ mice (bottom panels) across different timepoints. White arrows show targeted cells.
